# Hepatic Deletion of X-box Binding Protein 1 in Farnesoid X Receptor Null Mice Leads to Enhanced Liver Injury

**DOI:** 10.1101/2022.03.04.482879

**Authors:** Xiaoying Liu, Mahmoud Khalafalla, Chuhan Chung, Yevgeniy Gindin, Susan Hubchak, Brian LeCuyer, Alyssa Kriegermeier, Danny Zhang, Wei Qiu, Xianzhong Ding, Deyu Fang, Richard Green

**Author notes:** To whom correspondence should be addressed: Xiaoying Liu, PhD, Division of Gastroenterology and Hepatology, Department of Medicine, Northwestern University, McGaw Pavilion M-319, 240 E Huron Street, Chicago, IL 60611.

## Abstract

**Background & Aims:** Farnesoid X receptor (FXR) regulates bile acid metabolism and FXR null (*Fxr*^-/-^) mice have elevated bile acid levels and progressive liver injury. The inositol-requiring enzyme 1α (IRE1α)/X-box binding protein 1 (XBP1) pathway is a protective pathway of the unfolded protein response (UPR) that is activated in response to ER stress. In this study we sought to determine the role of the UPR in *Fxr*^-/-^ mice.

**Approach & Results:** We examined hepatic UPR gene and protein expression in 10- and 24-week-old wild type (WT) and *Fxr*^-/-^ mice. Hepatic XBP1 and other UPR pathways were activated in 24-week-old *Fxr*^-/-^ mice, but not WT mice. To further determine the role of the liver UPR activation in *Fxr*^-/-^ mice, we generated mice with FXR and liver-specific XBP1 double knockout (DKO, *Fxr*^-/-^*Xbp1*^LKO^) and *Fxr*^-/-^ *Xbp1*^fl/fl^ single knockout (SKO) mice and characterized their phenotypes at different ages. DKO mice demonstrated enhanced liver injury, apoptosis and fibrosis compared with SKO mice. RNA-seq revealed increased gene expression in apoptosis, inflammation and cell proliferation pathways in DKO mice. The proapoptotic C/EBP-homologous protein (CHOP) pathway was activated in DKO mice. At age 60 weeks, all DKO mice and no SKO mice spontaneously developed liver tumors.

**Conclusions:** The hepatic XBP1 pathway is activated in older *Fxr*^-/-^ mice and has a protective role. The potential interaction between XBP1 and FXR signaling may be important in modulating the hepatocellular cholestatic stress responses.

## Introduction

Farnesoid X receptor (FXR) is a bile acid-activated transcription factor that plays a central role in bile acid metabolism by regulating the expression of genes that are involved in bile acid synthesis and hepatocellular transport (1). Mice with FXR deletion have elevated serum bile acid levels and total bile acid pool size, and are susceptible to bile acid-induced cholestatic liver injury due to an attenuated ability to downregulate bile acid synthesis and excrete bile acids (2, 3). FXR null mice develop progressive liver inflammation and fibrosis as they age, and spontaneously develop liver tumors by 15 months of age (3–5). In contrast, FXR agonists may restore bile acid homeostasis and provide benefit to patients with cholestatic liver diseases. The FXR agonist obeticholic acid has been FDA-approved for primary biliary cholangitis (PBC), and FXR agonists are currently in clinical trials for the treatment of other cholestatic and metabolic liver diseases (6).

Endoplasmic reticulum (ER) stress occurs when ER homeostasis is perturbed due to excessive accumulation of unfolded/misfolded proteins. The liver is prone to ER stress given its large requirement for protein synthesis and folding to support its many metabolic and secretory functions. The unfolded protein response (UPR), comprising of inositol-requiring enzyme 1α (IRE1α), protein kinase R-like ER kinase (PERK) and activating transcription factor 6 (ATF6) signaling pathways, is a protective cellular response activated in response to ER stress. The hepatic UPR is important in the pathogenesis of several hepatic diseases, including viral hepatitis, non-alcoholic fatty liver disease, alpha-1 antitrypsin deficiency, alcohol-induced liver disease and ischemia-reperfusion injury (7, 8). The IRE1α/X-box binding protein 1 (XBP1) pathway is evolutionarily conserved, being present in both yeast and mammals. Phosphorylated and activated IRE1α causes an atypical splicing of XBP1 mRNA, resulting in the production of transcriptionally active XBP1 spliced (XBP1s) that regulates downstream target genes such as the chaperone ERdj4 to assist protein folding and EDEM which is involved in ER-associated degradation (ERAD) of accumulated proteins (9). Whole body XBP1 null mice are embryonically lethal (10) and liver-specific XBP1 deficient mice are unable to adequately resolve pharmacologically induced hepatic ER stress (11). Mice with hepatic XBP1 deletion also demonstrate decreased serum cholesterol, triglycerides and total bile acid pool size (12, 13).

ER stress and an inadequate compensatory UPR activation have been implicated in many liver diseases, including cholestatic liver disorders (14, 15). ER stress downregulates FXR expression in the liver and FXR inhibits ER stress-induced NLRP3 inflammasome activation (16). Thus, an inadequate UPR may lead to sustained ER stress, reduced FXR expression and accentuated inflammatory responses. We have previously shown that FXR agonists induce hepatic XBP1s activation, and hepatic XBP1s is induced in bile acid feeding and bile duct ligation models of cholestasis (17). However, the crosstalk between liver UPR and FXR signaling, especially in the setting of cholestatic liver injury, remains unclear. In this study, we developed novel FXR null mice with liver-specific deletion of XBP1 and utilized these mice to investigate the role of the UPR in bile acid injury.

### Experimental Procedures

#### Animal use and treatment

C57BL/6J wildtype (WT) and whole body FXR knockout (*Fxr*^-/-^) mice were obtained from Jackson Laboratory and colonies were established. Liver-specific XBP1 knockout (*Xbp1*^LKO^) and *Xbp1*^fl/fl^ colonies of mice in a C57BL/6J background have been maintained as described previously (18). *Fxr*^-/-^*Xbp1*^LKO^ double knockout (DKO) and *Fxr*^-/-^*Xbp1*^fl/fl^ single knockout (SKO) mice were generated by crossing *Fxr*^-/-^ mice with *Xbp1*^LKO^ and *Xbp1*^fl/fl^ mice. All the mice were housed on a 14-h light, 10-h dark cycle with free access to normal chow and water. Male mice were fasted for 4 h prior to sacrifice at ages of 8-10 weeks (labeled as 10-week for simplification), 24, 40 or 60 weeks. Blood was obtained by cardiac puncture, and the liver was removed and rinsed with ice-cold PBS and either immediately fixed in formalin or snap-frozen in liquid nitrogen. For total bile acid pool analysis, the liver, gallbladder and small intestine were collected from nonfasted male mice and immediately minced in 100% methanol. All protocols and procedures were approved by the Northwestern University Institutional Animal Care and Use Committee.

#### Serum biochemical analysis

Serum alanine aminotransferase (ALT) was measured by using a spectrophotometric assay according to the manufacturer’s protocol (Teco Diagnostics, Anaheim, CA).

#### Histology

Livers were fixed in 10% neutral buffered formalin, paraffin embedded, and sectioned. Immunohistological staining with hematoxylin and eosin (H&E), Sirius red, terminal deoxynucleotidyl transferase dUTP nick end labeling (TUNEL) and Ki67 was performed by Northwestern University Mouse Histology and Phenotyping Laboratory. The quantification of Sirius red was performed using ImageJ. The number of TUNEL stain positive cells and Ki67 positive cells were counted from 9-10 random fields per slide. The Ishak inflammation and fibrosis scores were assessed by 2 investigators (R.G. and A.K.) blinded to the study groups. The H&E staining at 60-week old was blindly assessed by a pathologist (X.D.).

#### Bile Acid Analysis

Serum bile acid levels were measured colorimetrically according to the manufacturer’s protocol (Genway Biotech, San Diego, CA). Enterohepatic total bile acid pool size and species were determined by HPLC using the method of Heumann as previously described (13). For hepatic bile acid analysis, 100 mg of livers were homogenized in 1 ml 70% ethanol. The homogenates were incubated at 60°C for 2 hours and centrifuged at 4000 rpm for 10 minutes. 100 μl of the supernatant was dried down, resuspended in 100 μl PBS and used to determine bile acid concentration spectrophotometrically.

#### RNA Extraction and quantitative PCR

Total RNA was extracted from frozen livers using Trizol according to the manufacturer’s protocol (Invitrogen Life Technologies, Carlsbad, CA) and cDNA was made with qScript cDNA synthesis kit (Quanta Bioscience, Gaithersburg, MD). Quantitative PCR (qPCR) was performed as described previously (19) and all primers were synthesized by Sigma (St. Louis, MO).

#### RNA-seq

Stranded mRNA-seq was conducted in the Northwestern University NUSeq Core Facility. Briefly, total RNA examples were checked for quality using RINs generated from Agilent Bioanalyzer 2100. RNA quantity was determined with Qubit fluorometer. The Illumina TruSeq Stranded mRNA Library Preparation Kit was used to prepare sequencing libraries from 1 μg of high-quality RNA samples (RIN>7) using the methodology of manufacturer. This procedure includes mRNA purification and fragmentation, cDNA synthesis, 3’ end adenylation, Illumina adapter ligation, library PCR amplification and validation. lllumina HiSeq 4000 sequencer was used to sequence the libraries with the production of single-end, 50 bp reads at the depth of 20-25 M reads per sample. Gene expression was quantified from RNA-seq reads with Salmon (20) using mouse reference genome version GRCm38 obtained from GENCODE (21). Read counts were converted to counts-per-million using edgeR (22). Pathway enrichment analysis was accomplished by calculating a normalized enrichment score against Molecular Signatures Database hallmark gene set collection (23) as implemented in clusterProfiler (24).

#### Western blotting

Protein extraction was made from frozen liver tissues using T-Per protein extraction reagent (Thermo Scientific, Waltham, MA) with protease inhibitors cocktail (MilliporeSigma, Burlington, MA) and phosphatase inhibitors (Thermo Scientific, Waltham, MA). Protein quantification and immunoblotting were performed as described previously (19).

#### Statistics

Data are shown as means ± SEM and graphed with Prism (GraphPad, San Diego, CA). Comparison between two groups was performed by two-tailed Student’s t-test. Statistical significance was defined as P values of less than 0.05.

## Results

### Hepatic IRE1α/XBP1 pathway was activated in 24-week-old *Fxr*^-/-^ mice

*Fxr*^-/-^ mice develop spontaneous, progressive liver injury and fibrosis (4). To establish whether there was UPR activation indicative of ER stress in *Fxr*^-/-^ mice as they age, we first examined the activation of the three UPR pathways in the livers of WT and *Fxr*^-/-^ mice at 10 weeks and 24 weeks of age. As shown in Figure 1A and Supplementary Figure S1, the nuclear protein expression of XBP1s was significantly upregulated in 24-week-old *Fxr*^-/-^ mice compared to 10-week-old *Fxr*^-/-^ mice. The mRNA expression of *Xbp1s* and its downstream target genes *ERdj4* and *Edem* was significantly higher in 24-week-old *Fxr*^-/-^ mice compared to 10-week-old *Fxr*^-/-^ mice (Figure 1B). The protein expression of the XBP1s upstream activator phosphorylated IRE1α (p-IRE1α) also significantly increased in 24-week-old *Fxr*^-/-^ mice (Figures 1A and Supplementary Figure S1). In contrast, there was no significant change in hepatic nuclear XBP1s protein level, mRNA expression of *Xbp1s*, *ERdj4* and *Edem* and IRE1α phosphorylation when comparing 10-week-old vs. 24-week-old WT mice (Figures 1A, 1B and Supplementary Figure S1). Hepatic *Ire1*α mRNA expression did not change, although total IRE1α protein expression was reduced in 24-week-old mice in both genotypes (Figure 1A, 1B and Supplementary Figure S1). These data demonstrate that the hepatic IRE1α/XBP1 pathway was activated in 24-week-old compared to 10- week-old *Fxr*^-/-^ mice, but not in WT mice of similar ages.

**Figure 1.**
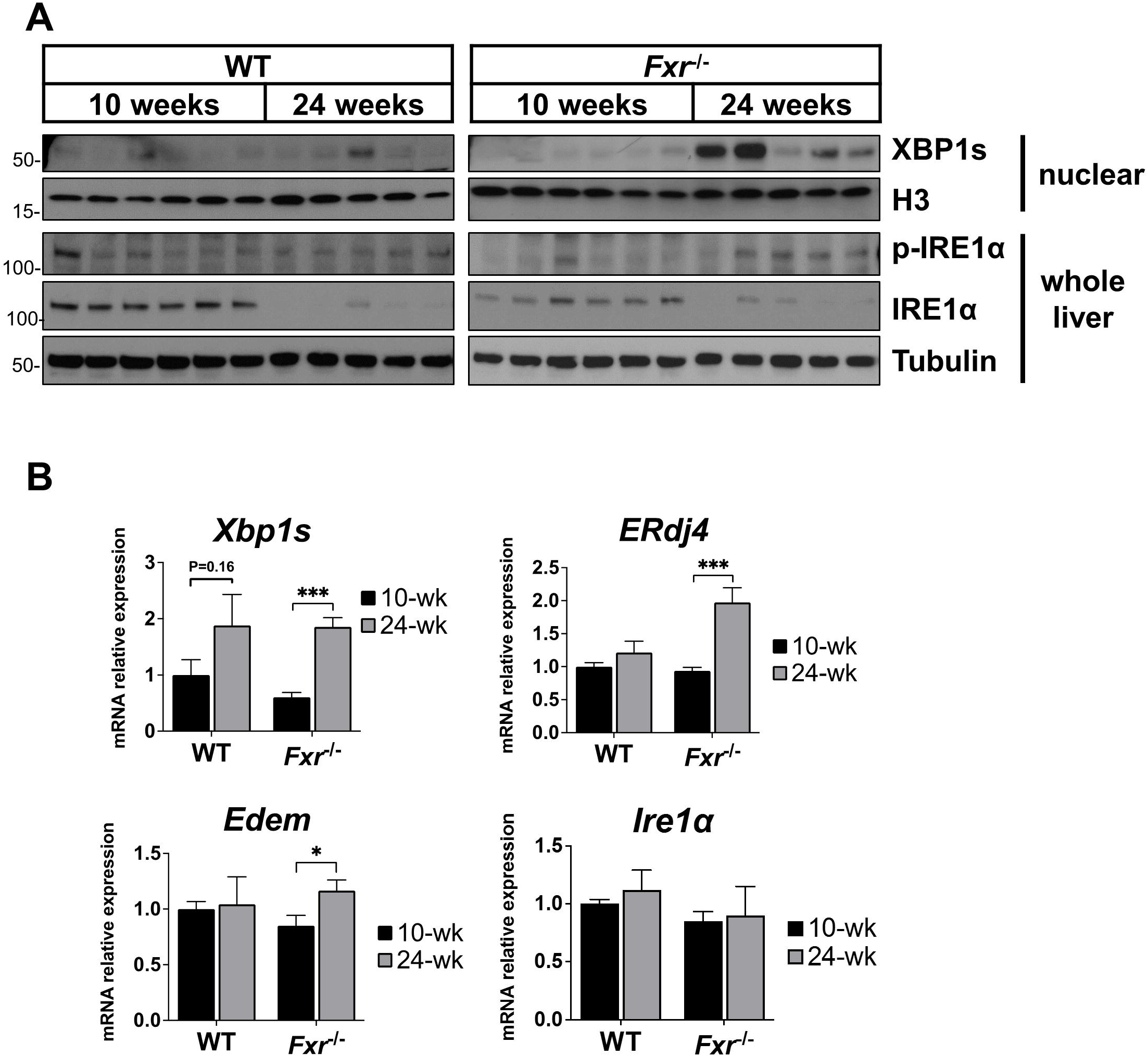
Hepatic IRE1α/XBP1 pathway was activated in 24-week-old *Fxr*^-/-^ mice. The hepatic expression of the IRE1α/XBP1 pathway was examined in male 10-week-old (n=6) and 24-week-old (n=5) WT and *Fxr*^-/-^ mice. (A) Nuclear XBP1s, whole liver p-IRE1α and IRE1α protein expression was examined by western blotting with histone 3 (H3) and tubulin as loading controls for nuclear and whole liver lysates, respectively. (B) Hepatic gene expression of *Xbp1s*, *ERdj4*, *Edem* and *Ire1*α was measured by qPCR. *P<0.05, ***P<0.001.

The PERK pathway of the UPR is activated through auto-phosphorylation of PERK, which then phosphorylates downstream eukaryotic initiation factor 2α (eIF2α) and inhibits global protein translation. Phosphorylated-eIF2α (p-eIF2α) also selectively increases translation of a number of mRNAs, such as activating transcription factor 4 (ATF4). There was a strong trend towards increased phosphorylated-PERK (p-PERK) expression in 24-week-old *Fxr*^-/-^ mice compared with 10-week-old *Fxr*^-/-^ mice (P=0.06), while WT mice demonstrated similar levels (Figure 2A, Supplemental Figure S2). Hepatic expression of p-eIF2α was significantly lower in 24-week-old compared to 10-week-old mice in both genotypes. PERK and eIF2α protein expression were also lower in 24-week-old mice compared to 10-week-old mice. The protein expression of ATF4 increased in 24-week-old *Fxr*^-/-^ mice compared with 10-week-old *Fxr*^-/-^ mice, but not in WT mice (Figure 2A, Supplemental Figure S2). The nuclear protein expression of ATF4 downstream target, C/EBP-homologous protein (CHOP) was also increased in 24-week-old *Fxr*^-/-^ mice compared to 10-week-old *Fxr*^-/-^ mice, but not in WT mice (Figure 2A, Supplemental Figure S2). Consistently, the gene expression of the CHOP downstream target, death receptor 5 (*Dr5*) significantly increased in 24-week-old *Fxr*^-/-^ mice compared with 10-week-old *Fxr*^-/-^ mice, but not in WT mice (Figure 2B). The hepatic gene expression of the ATF6 pathway targets, *Bip* and *Hyou1*, was also significantly higher in 24-week-old *Fxr*^-/-^ mice compared with 10-week-old *Fxr*^-/-^ mice, but not in WT mice (Figure 2C). Together these data indicate that ER stress and UPR activation occurred in older *Fxr*^-/-^, but not WT mice.

**Figure 2.**
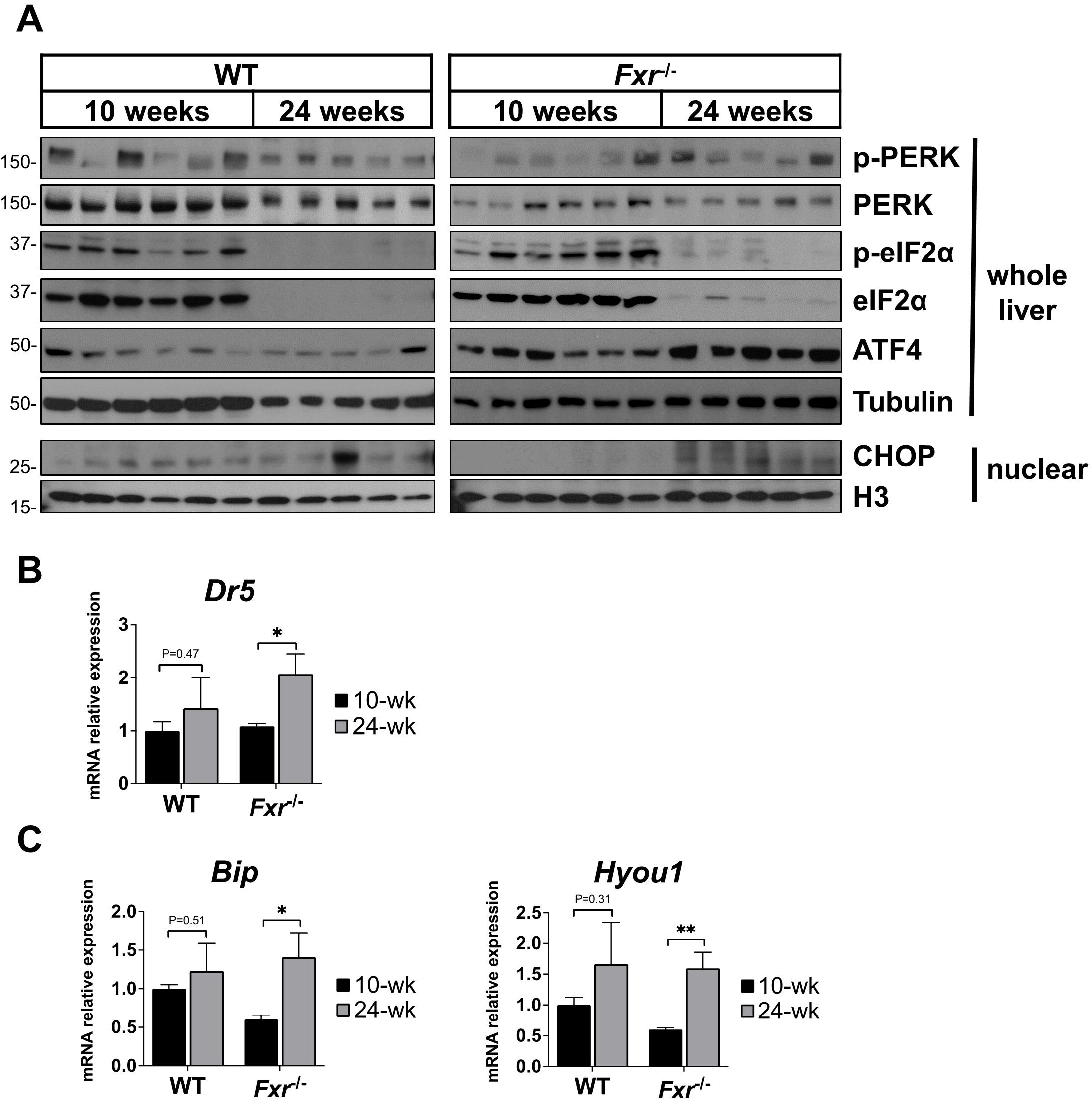
Other hepatic UPR pathways were activated in 24-week-old *Fxr*^-/-^ mice. The hepatic expression of the PERK and ATF6 pathway genes and proteins was examined in male 10-week-old (n=6) and 24-week-old (n=5) WT and *Fxr*^-/-^ mice. (A) The protein expression of whole liver p-PERK, PERK, p-eIF2α, eIF2α, ATF4 and nuclear CHOP was examined by western blotting. H3 and tubulin were used as loading controls for nuclear and whole liver lysates, respectively. Hepatic gene expression of (B) CHOP downstream target *Dr5* and (C) ATF6 downstream targets *Bip* and *Hyou1* was measured by qPCR. *P<0.05, **P<0.01.

### Hepatic deficiency of XBP1 promoted liver injury and fibrosis in *Fxr*^-/-^ mice at 10 weeks old

To determine the role of hepatic XBP1 activation in *Fxr*^-/-^ mice, we developed colonies of *Fxr*^-/-^/liver-specific XBP1KO double knockout (DKO, *Fxr*^-/-^*Xbp1*^LKO^) and control *Fxr*^-/-^/XBP1-flox single knockout (SKO, *Fxr*^-/-^*Xbp1*^fl/fl^) mice (Supplementary Figure S3), and characterized their phenotypes at 10 weeks of age. DKO mice had 1.5-fold higher serum ALT levels compared to SKO mice, being 209.1±16.6 *vs* 83.1±17.2 U/L, P<0.001 (Figure 3A). H&E staining showed mild inflammation with a strong trend towards higher Ishak inflammation scores in DKO mice (P=0.06) (Figures 3B, 3C). DKO mice had significantly greater liver fibrosis compared to SKO mice as evidenced by increased Sirius red staining and higher Ishak fibrosis scores (Figures 3B, 3D, 3E). In addition, more TUNEL stain positive cells were observed in DKO mice livers compared to SKO mice (Figures 3B and 3F), consistent with greater liver injury in DKO mice. These data indicate that the absence of liver XBP1 in *Fxr*^-/-^ mice worsens tissue injury and increases fibrosis.

**Figure 3.**
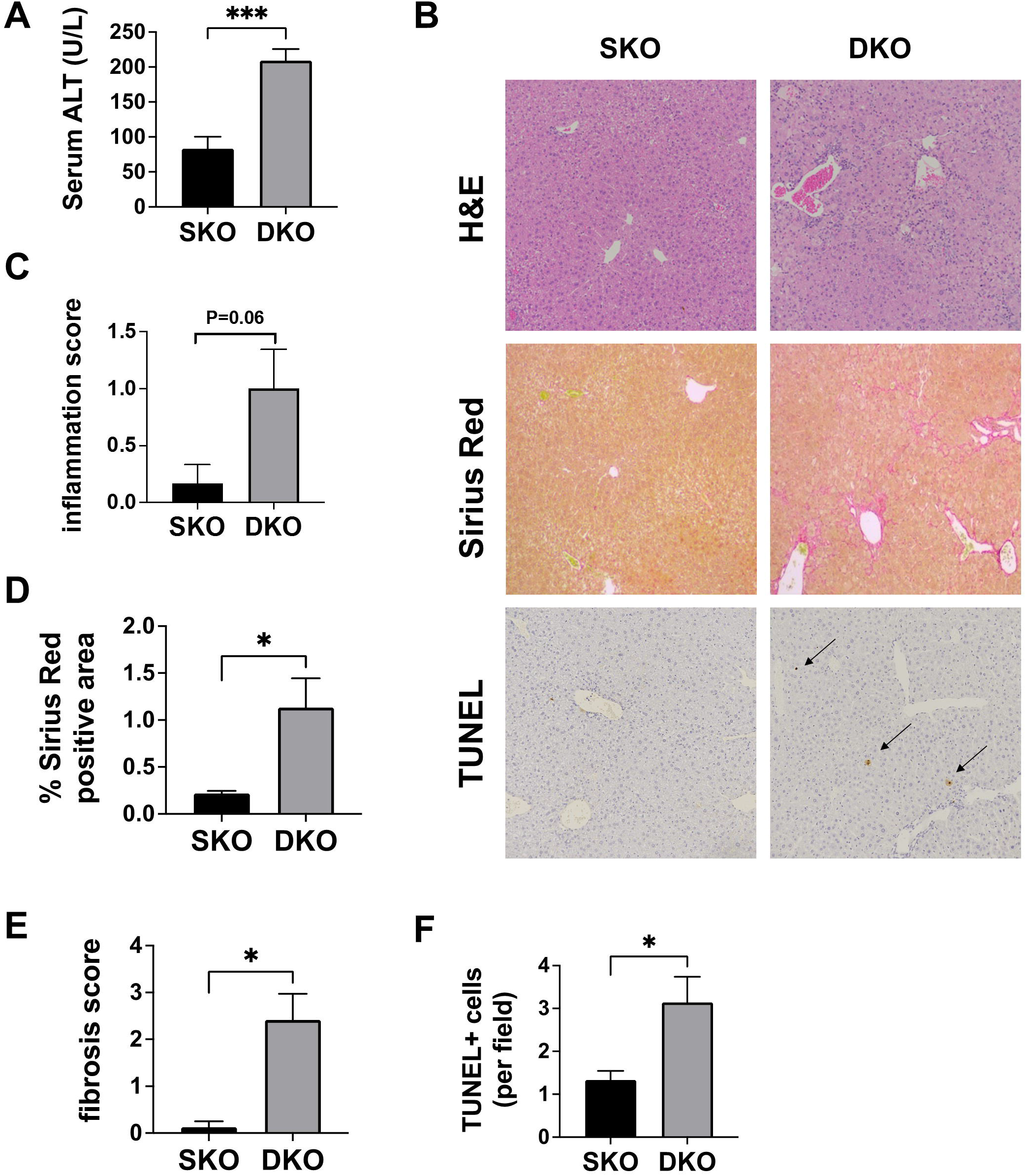
*Fxr*^-/-^*Xbp1*^LKO^ mice had greater liver injury and fibrosis compared to *Fxr*^-/-^*Xbp1*^fl/fl^ mice at 10 weeks old. The phenotypes of 10-week-old male *Fxr*^-/-^*Xbp1*^fl/fl^ mice (SKO, n=4) and *Fxr*^-/-^*Xbp1*^LKO^ (DKO, n=6) mice were characterized. (A) Serum ALT levels. (B) Representative images of H&E, Sirius red and TUNEL staining (arrows) (200x magnification). (C) Ishak inflammation scores. (D) Percentage of Sirius red stain positive area per field. (E) Ishak fibrosis scores. (F) Number of TUNEL stain positive cells per field. *P<0.05, ***P<0.001.

### RNA-seq demonstrated increased gene expression in inflammation, apoptosis and proliferation pathways in DKO mice

In order to delineate the mechanisms by which hepatic *Xbp1* deletion promotes liver injury and fibrosis, we performed RNA-seq on 10-week-old DKO and SKO mice livers. The first principal component of the RNA-seq data accounted for 76% of variance in hepatic gene expression (Figure 4A). Gene set enrichment analysis using Hallmark gene set collection revealed that multiple immune-related pathways (inflammatory_response, TNFα_signaling_via_NFkb, IL6_JAK_STAT3_signaling and IL2_STAT5_signaling) were upregulated in DKO mice compared to SKO mice (Figure 4B). The Apoptosis pathway was also upregulated in DKO mice. Cell proliferation related (G2M_checkpoint, E2F_targets, p53_pathway and Myc_target_v2) and angiogenesis pathways were similarly upregulated in DKO mice compared to SKO mice. These results were consistent with the more injurious phenotype of DKO mice. Bile acid metabolism and fatty acid metabolism pathways were downregulated in DKO mice, which is consistent with previous reports in hepatic-specific XBP1 knockout mice (13). Therefore, the transcriptome profile of the DKO mice reflects the enhanced tissue injury that occurs in the absence of a functional XBP1 pathway in *Fxr*^-/-^ mice.

**Figure 4.**
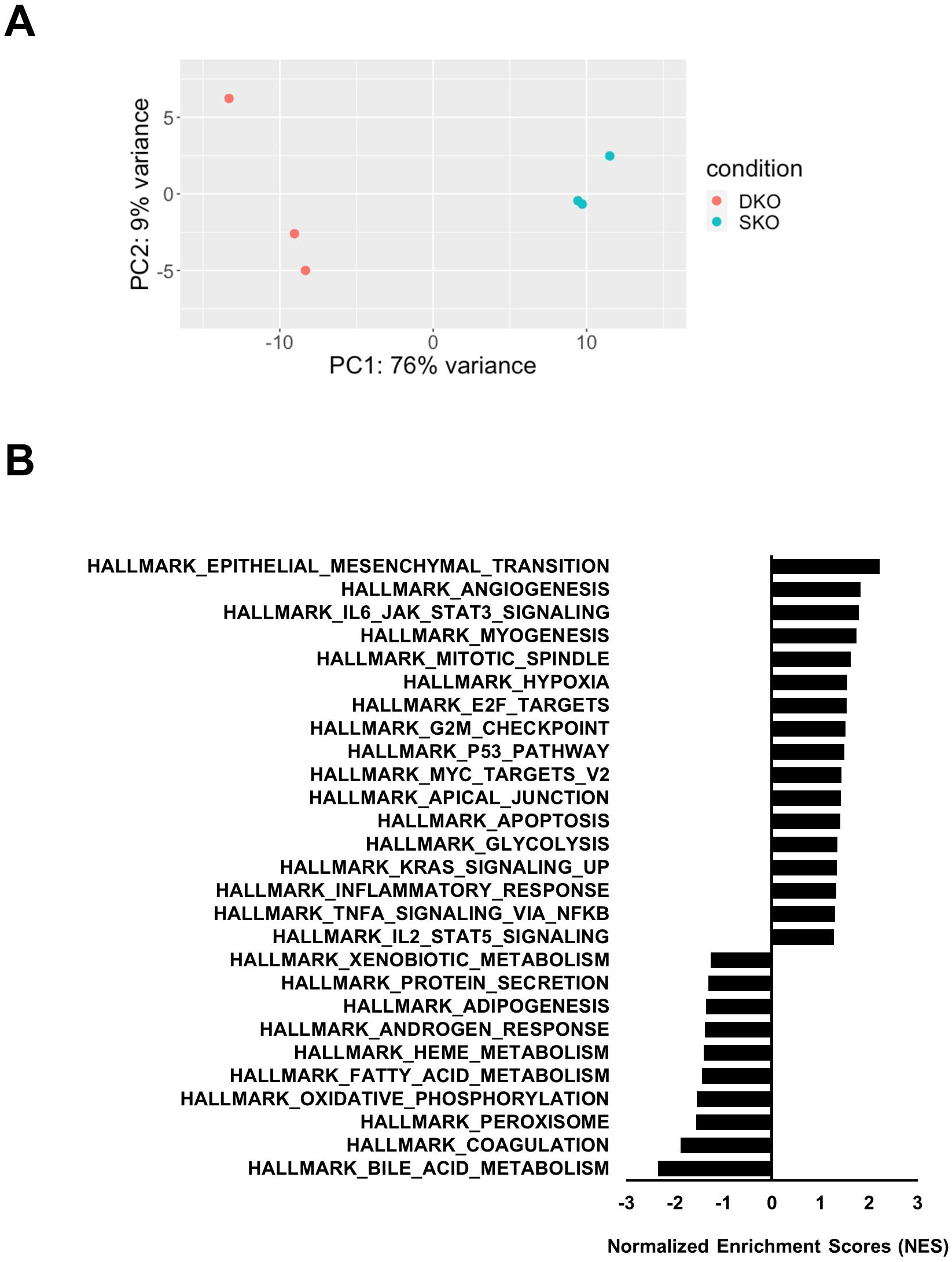
RNA-seq revealed that *Fxr*^-/-^*Xbp1*^LKO^ mice had increased hepatic expression of apoptosis, inflammation and proliferation pathways. Hepatic transcriptome profiles of 10-week-old male *Fxr*^-/-^*Xbp1*^fl/fl^ mice (SKO, n=3) and *Fxr*^-/-^*Xbp1*^LKO^ mice (DKO, n=3) were examined by RNA-seq. (A) PCA plot demonstrating the segregation and clustering of the two groups. (B) Gene set enrichment analysis using the Hallmark collection showing significantly enriched pathways with a P-adj<0.05.

### The hepatic eIF2α/ATF4/CHOP pathway was activated in DKO mice

The eIF2α/ATF4/CHOP pathway mediates ER stress-induced cell apoptosis, and the protein expression of p-eIF2α and ATF4 was significantly higher in 10-week-old DKO mice compared to SKO mice (Figure 5A, Supplementary Figure S4). Gene expression of *Chop* and its downstream target *Dr5* was 3-fold (P<0.05) and 5-fold (P<0.01) higher in DKO compared to SKO mice, respectively (Figure 5B). The proapoptotic protein BAX was also upregulated in DKO compared to SKO mice (Figure 5A). Protein expression of p-PERK was similar in both genotypes, although PERK expression was lower in DKO mice (Supplementary Figure S4). To evaluate if these changes were due, at least in part, to hepatic XBP1 deletion, we examined the activation of the eIF2α/ATF4/CHOP pathway in *Xbp1*^LKO^ and *Xbp1*^fl/fl^ mice. *Xbp1*^LKO^ mice had higher hepatic expression of p-eIF2α, *Chop* and *Dr5* compared to *Xbp1*^fl/fl^ mice (Figures 5A, 5B), similar to the findings in DKO mice compared to SKO mice. In contrast, the marked increases in ATF4 and BAX protein expression was not observed comparing *Xbp1*^LKO^ to *Xbp1*^fl/fl^ mice.

**Figure 5.**
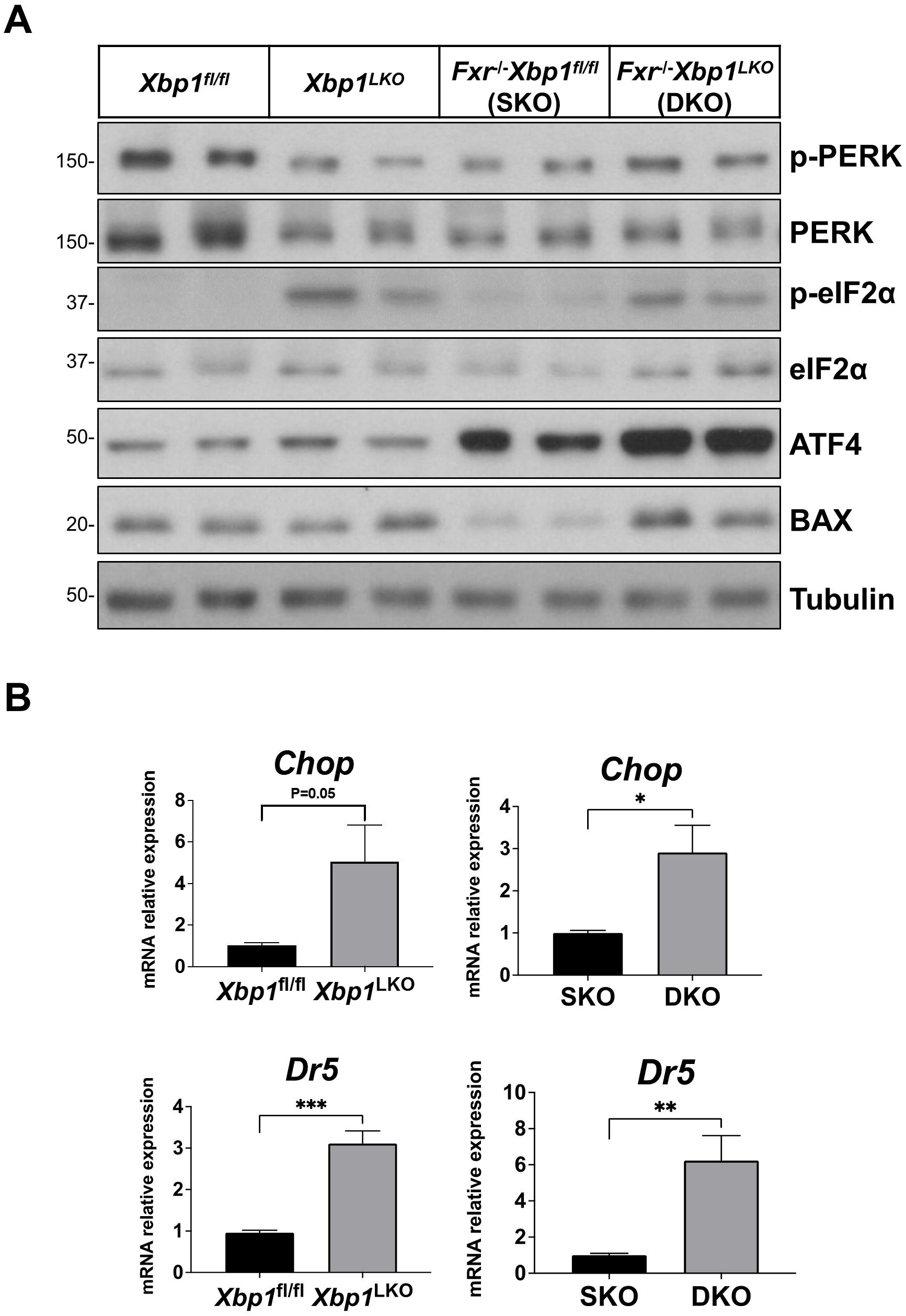
The hepatic expression of the eIF2α/ATF4/CHOP pathway was increased in *Fxr Xbp1*^LKO^ mice. Liver protein and RNA were isolated from 10-week-old male *Xbp1*^fl/fl^ (n=4), *Xbp1*^LKO^ (n=4), *Fxr*^-/-^*Xbp1*^fl/fl^ (SKO, n=4) and *Fxr*^-/-^*Xbp1*^LKO^ mice (DKO, n=6). (A) Western blotting examining p-PERK, PERK, p-eIF2α, eIF2α, ATF4 and BAX protein expression in whole liver. Tubulin was used as a loading control. Each lane represents a pooled sample of 2-3. (B) qPCR analysis of *Chop* and *Dr5*. *P<0.05, **P<0.01, ***P<0.001.

### SKO and DKO mice had similar total bile acid pool size, but differing bile acid composition

We have previously shown that hepatic XBP1 deficiency modulates bile acid metabolism (13), therefore we investigated the bile acid content in DKO and SKO mice. The total bile acid pool size was similar in DKO and SKO mice (65.3±4.6 vs. 52.2±6.2 μmol/100g mouse, respectively, P=0.27) (Figure 6A). The percentage of taurocholic acid (TCA) increased and the percentage of tauromuricholic acid (TMCA) decreased significantly in DKO compared with SKO mice (70.4±1.0 % vs 55±1.7 % for TCA; 27.2±1.0 % vs 41.9±2.4 % for TMCA; P<0.01) (Figure 6B). Consistent with the change of bile acid composition, the hydrophobicity index was higher in DKO compared with SKO mice (−0.21±0.01 vs −0.33±0.03, P<0.05, Figure 6C), while the serum and hepatic bile acid levels did not differ (Figures 6D, 6E). Despite the similar bile acid pool size, hepatic protein expression of the bile acid synthesis enzyme CYP7A1 was higher while the bile acid transporters *Bsep* and *Ntcp* gene expression was lower in DKO compared with SKO mice (Figures 6F, 6G). These findings indicate that loss of liver XBP1 in addition to FXR deficiency further disrupts multiple genes involved in bile acid synthesis and homeostasis.

**Figure 6.**
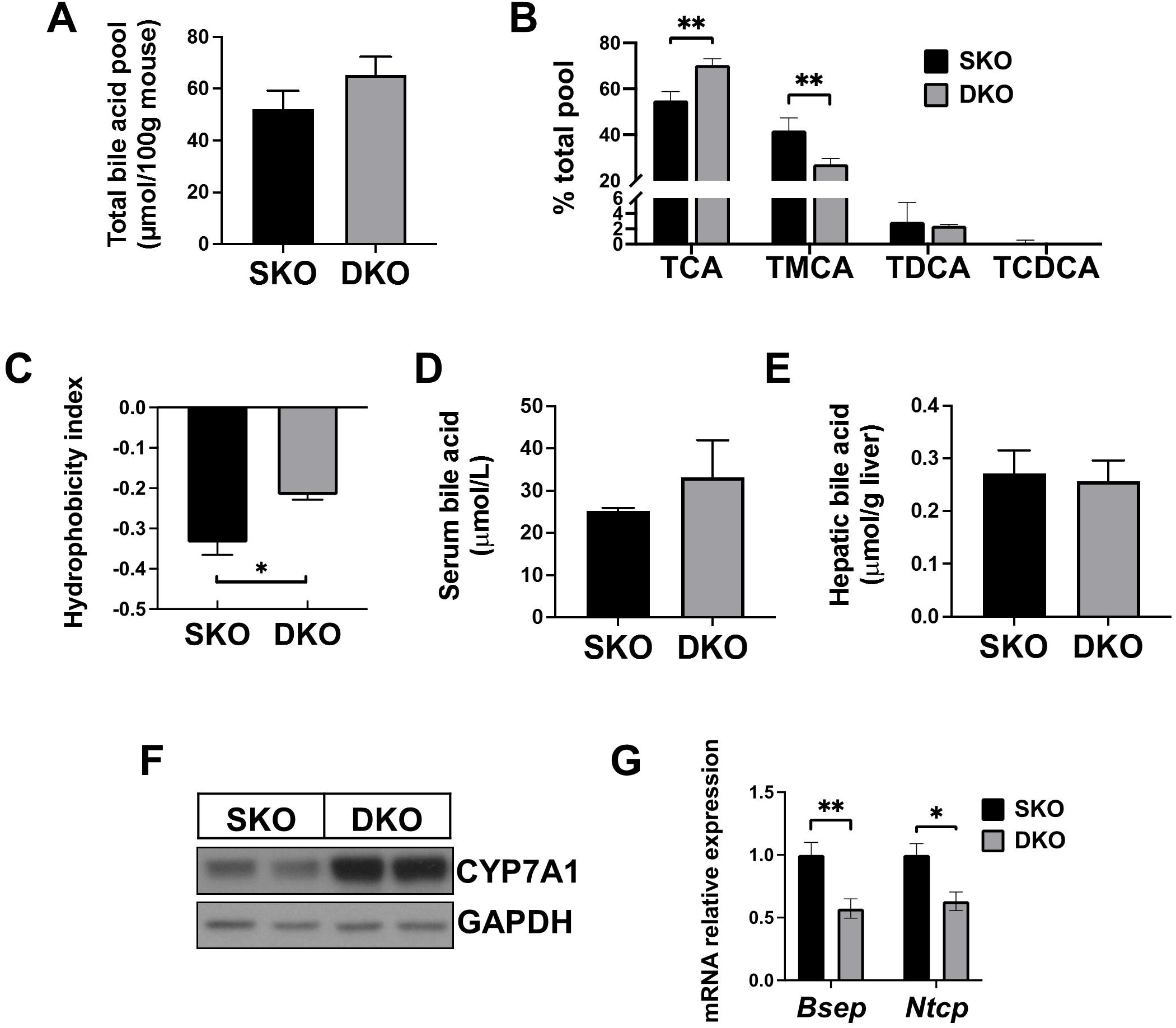
The bile acid profiles in *Fxr*^-/-^*Xbp1*^fl/fl^ and *Fxr*^-/-^*Xbp1*^LKO^ mice. (A) Total bile acid pool size and (B) bile acid composition were measured by HPLC in male *Fxr*^-/-^*Xbp1*^fl/fl^ mice (SKO, n=4) and *Fxr*^-/-^ *Xbp1*^LKO^ (DKO, n=3) mice. (C) Bile acid pool hydrophobicity index. (D) Serum bile acid levels were measured (SKO, n=3; DKO, n=4). (E) Hepatic bile acids levels were measured (SKO, n=6; DKO, n=6). (F) Western blotting demonstrated increased hepatic CYP7A1 protein expression in DKO mice compared to SKO mice. Each lane represents a pooled sample of 2-3. (G) Hepatic gene expression of *Bsep* and *Ntcp* was measured by qPCR (SKO, n=4; DKO, n=6). *P<0.05, **P<0.01.

### DKO mice had greater liver injury and fibrosis compared to SKO mice at 24 and 40 weeks of age

In order to determine whether the enhanced liver injury in DKO mice persists at older ages, we subsequently compared DKO and SKO mice at 24 and 40 weeks of age. H&E staining and Ishak scores indicated increased liver inflammation in DKO mice compared to SKO at 40-week-old, without a genotype-specific difference at 24 weeks (Figures 7A–C). Serum ALT levels remained significantly higher in DKO compared with SKO mice at 24-week-old (106.7±8.7 *vs* 63.8±3.5 U/L, P<0.01); but was similar between the two groups at 40-week-old (127.2±13.6 *vs* 92.8±9.9 U/L, P=0.08) (Figures 7B, 7C). At age 24 and 40 weeks, DKO mice had more fibrosis than SKO mice, evident by Sirius red staining quantification and Ishak fibrosis scores (Figures 7A-C). Thus, the absence of hepatic XBP1 leads to persistent liver injury in the DKO mice.

**Figure 7.**
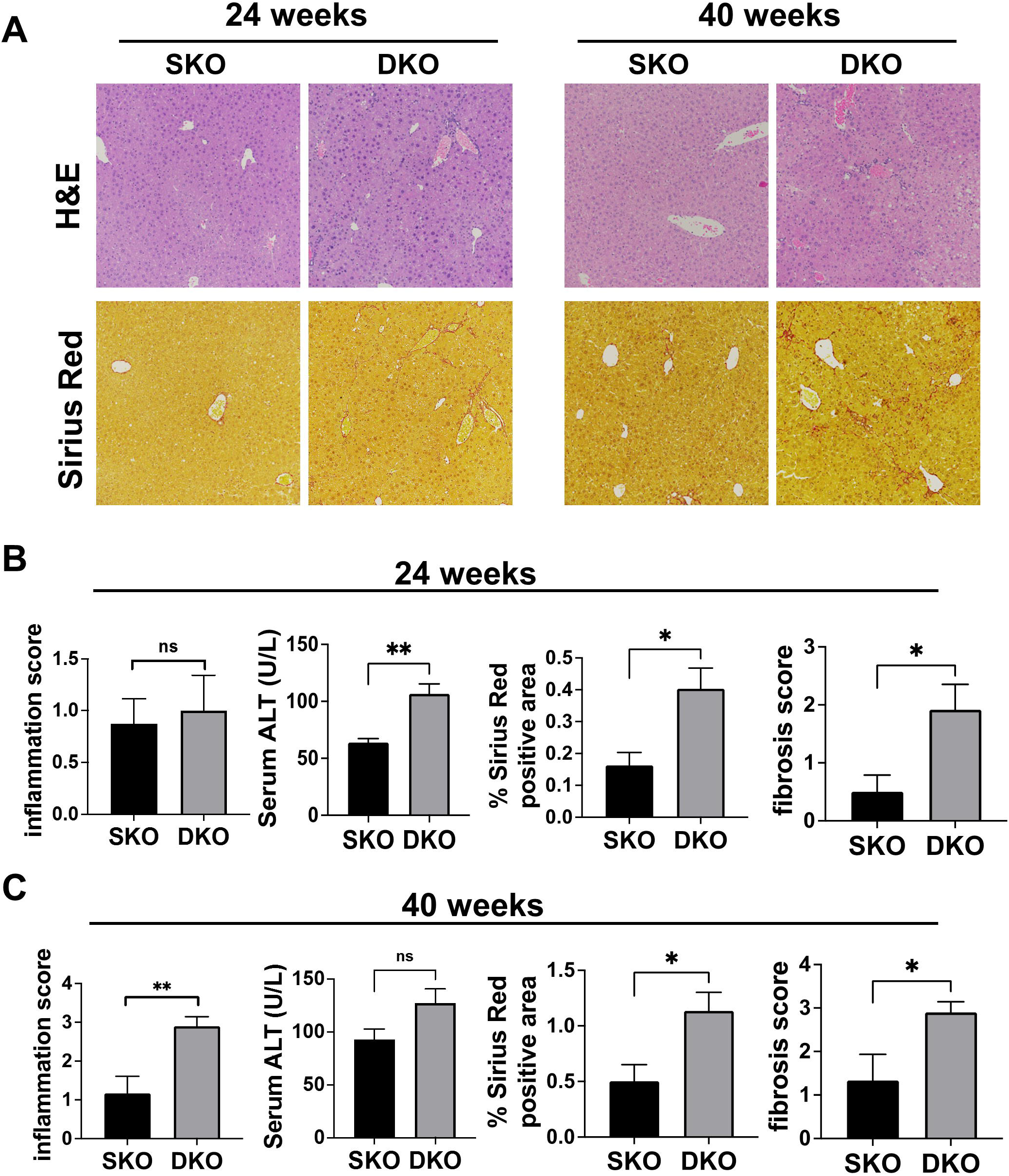
The phenotypes of *Fxr*^-/-^*Xbp1*^fl/fl^ and *Fxr*^-/-^*Xbp1*^LKO^ mice at 24 and 40 weeks of age. (A) Representative images of H&E and Sirius red staining from 24-week-old and 40-week-old male *Fxr*^-/-^ *Xbp1*^fl/fl^ mice (SKO) and *Fxr*^-/-^*Xbp1*^LKO^ (DKO) mice (200x magnification). (B) Ishak inflammation scores, serum ALT, percentage of Sirius red stain positive area and Ishak fibrosis scores of 24-week-old male SKO mice (n=4-6) and DKO mice (n=6). (C) Ishak inflammation scores, serum ALT, percentage of Sirius red stain positive area and Ishak fibrosis scores of 40-week-old male SKO mice (n=3-5) and DKO mice (n=5-6). *P<0.05, **P<0.01.

### Hepatic deficiency of XBP1 promoted liver tumorigenesis in 60 weeks old *Fxr*^-/-^ mice

Since RNA-seq study revealed that multiple cell proliferation pathways were upregulated in DKO mice, we aged DKO and SKO mice to 60 weeks old to assess the effect of hepatic *Xbp1* deletion on tumor development in *Fxr*^-/-^ mice. We did not observe any liver tumor formation in *Fxr*^-/-^ mice (n=4), SKO mice (n=10) or *Xbp1*^LKO^ mice (n=7) at age 58-60 weeks. In contrast, 6 of 6 (100%) of DKO mice spontaneously developed multiple liver tumors at 60 weeks of age (Figure 8A). Tumor nodules showed diffuse hepatocellular proliferation with increased hepatocyte density, loss of normal hepatic architecture, and lack of portal tracts. Tumors exhibited marked nuclear crowding, increased nuclear-to-cytoplasmic ratio, dense nuclear chromatin, and increased proliferation index. The H&E histologic findings were consistent with hepatocellular adenomas. The liver to body weight ratio was significantly higher in DKO mice compared to SKO mice (Figure 8B). Ki67 staining showed greater than 30-fold increase in DKO mice compared to SKO mice (Figure 8A, 8C), while serum ALT was similar between the two groups (Figure 8D). No hepatic tumors were observed in DKO or SKO mice at 10, 24 or 40 weeks of age. The hepatic expression of the cell cycle protein Cyclin D1 was upregulated in DKO mice compared to SKO mice at 10 and 24 weeks of ages but was not significantly different at age 40 weeks (Figure 8E, Supplementary Figure S5). The hepatic Cyclin D1 protein expression was similar between *Xbp1*^LKO^ mice and *Xbp1*^fl/fl^ mice (Supplementary Figure S5).

**Figure 8.**
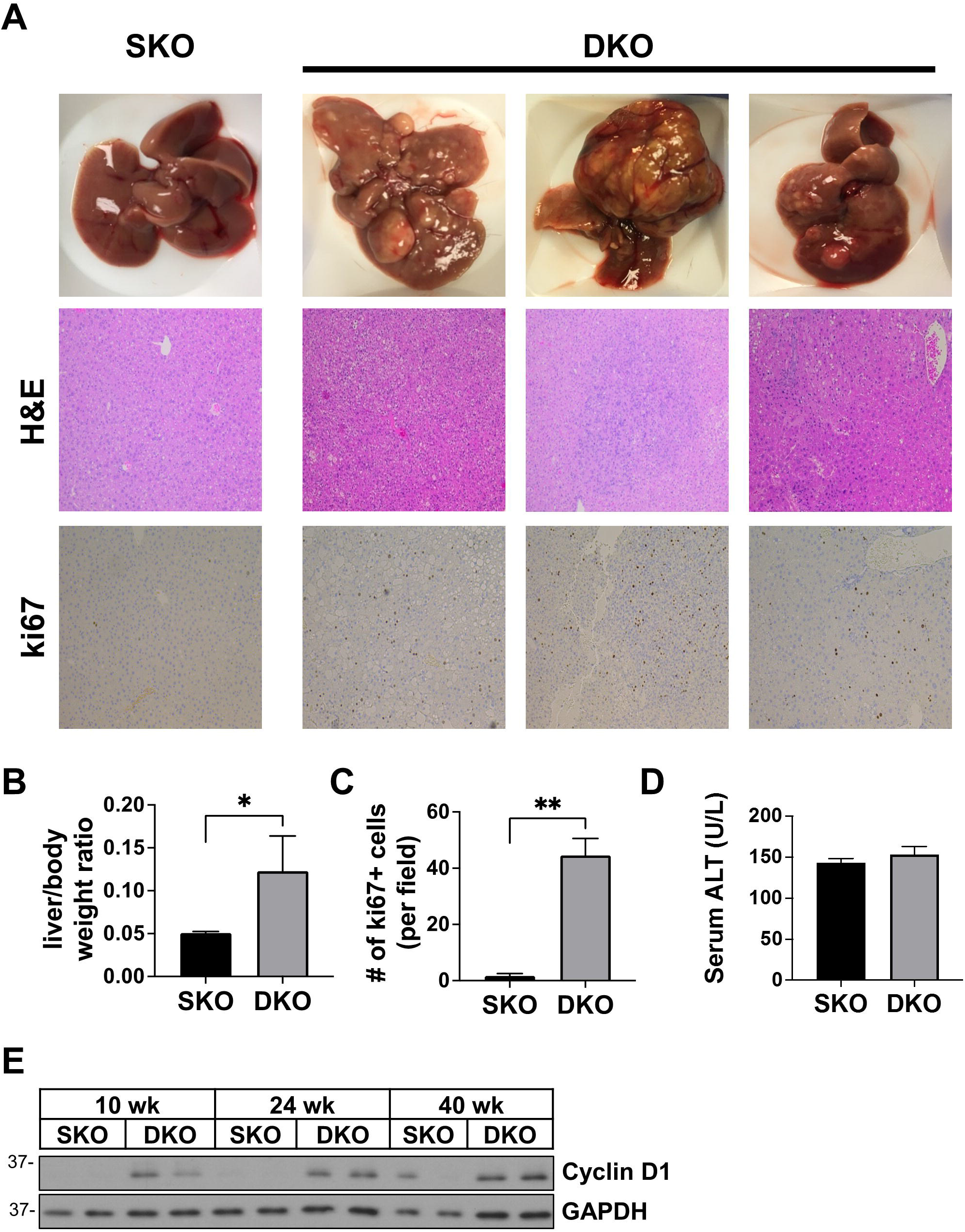
*Fxr*^-/-^*Xbp1*^LKO^ mice spontaneously developed liver tumors at 60 weeks of age. Chow-fed male *Fxr*^-/-^*Xbp1*^fl/fl^ mice (SKO, n=10) and *Fxr*^-/-^*Xbp1*^LKO^ (DKO, n=6) mice were aged to 60 weeks of age. (A) Representative images of gross liver, H&E and Ki67 staining (200x magnification). (B) Liver weight to body weight ratio. (C) The numbers of Ki67 positive cells per field. (D) Serum ALT. (E) Western blotting demonstrating hepatic Cyclin D1 protein expression in male SKO and DKO mice at 10-, 24- and 40-week-old. GAPDH was used as a loading control. Each lane represents a pooled sample of 2-3. *P<0.05, **P<0.01.

## Discussion

The nuclear receptor FXR plays a central role in hepatocellular physiology and pathophysiology by regulating several important hepatic metabolic and transporter processes, as well as eliciting cellular protective functions in immunologic and non-parenchymal cells. FXR null mice are a well characterized murine model of bile acid toxicity and cholestasis since intrahepatic bile acid concentrations are elevated. Decreased expression of FXR has been associated with aging and diseases such as PBC and HCC (25–27). It has been previously reported that FXR deficient mice develop spontaneous progressive inflammation, steatosis, fibrosis and HCC, which mimics human HCC progression (4). In this study, we initially demonstrated that the liver IRE1α/XBP1 and other UPR pathways were activated in 24-week-old *Fxr*^-/-^ mice. Increased hepatic bile acid concentrations are known to induce ER stress and activate the UPR (28, 29), and the ER stress that occurs in *Fxr*^-/-^ mice as they age is likely due to the persistently increased bile acid levels in these mice (5).

In order to study the specific role of hepatic XBP1 pathway activation in *Fxr*^-/-^ mice, we developed DKO mice with both global loss of FXR and hepatocyte-specific deletion of XBP1. DKO mice demonstrated significant liver injury and fibrosis as early as 10 weeks of age compared with SKO mice. This suggests that the enhanced hepatic XBP1 pathway is protective in FXR deficient mice, which is consistent with the known adaptive and protective role of XBP1 in reducing or resolving ER stress. A recent transcriptome study using liver biopsies from PSC patients demonstrated that the expression of UPR genes was lower in the group of patients who were at increased risk for adverse PSC-related clinical events (15). Furthermore, we have shown in murine models that weanling mice are unable to adequately activate the hepatic XBP1 and other UPR pathways in response to bile acid or pharmacologically-induced ER stress, and this inadequate hepatic XBP1 activation causes increased liver injury and apoptosis (30). These data together further demonstrate the importance of a functional XBP1 pathway and intact UPR in lessening cholestatic or other forms of liver injury. FXR agonists activate the hepatic XBP1 pathway in mice and *in vitro* (17). The potent FXR agonist obeticholic acid, that is FDA-approved for PBC, may exert its beneficial effects, at least in part, through activating the protective UPR pathway. Ursodeoxycholic acid (UDCA) is also approved for the treatment of PBC and tauro-UDCA is a chemical chaperone that reduces ER stress. Therefore, UDCA may be hepatoprotective, in part, due to its chemical chaperone properties in addition to its other known mechanisms of action.

Sustained ER stress can lead to cell death. Mice with hepatic XBP1 deletion that are treated with the pharmacologic ER stress inducer tunicamycin fail to adequately resolver ER stress and have resultant increased apoptosis (11). Consistent with these data, our hepatic TUNEL staining showed more apoptosis and RNA-seq demonstrated enhanced apoptosis pathway expression in DKO mice. CHOP and its downstream activation of DR5 induce apoptosis during ER stress (31) and the hepatic gene expression of *Chop* and *Dr5* was consistently higher in the DKO mice. This was likely due to, at least in part, to the increased liver p-eIF2α and ATF4 protein expression in DKO mice. The significantly increased hepatic protein expression of ATF4, BAX and Cyclin D1 in DKO mice compared to SKO mice, did not occur in *Xbp1*^LKO^ mice compared to *Xbp1*^fl/fl^ mice when FXR signaling was intact. These changes seen in DKO mice may be attributed to further increases of ER stress due to elevated bile acid levels or potential crosstalk between liver XBP1 and FXR signaling.

We have previously shown that hepatic XBP1 deletion decreases the total bile acid pool (13). The total bile acid pool size in our study was similar in the DKO and SKO mice indicating that the FXR-dependent and XBP1-dependent changes in bile acid metabolism likely both occur. The hepatic protein expression of the bile acid synthesis gene CYP7A1 was elevated in DKO mice compared to SKO mice, which may contribute to the equalization of bile acid pool size. Furthermore, the total bile acid pool was more hydrophilic and thus less toxic in DKO mice compared to SKO mice. Therefore, the adverse effects of hepatic XBP1 deletion in FXR deficient mice could not be attributed to more injurious bile acid levels or species. Previous studies have demonstrated a role of intestinal FXR signaling in regulating the microbiome, bile acid composition and thereby metabolic functions (32). Further investigations are needed to determine potential differences in gut microbiome in DKO and SKO mice.

Although it has been reported that *Fxr*^-/-^ mice spontaneously develop HCC by 15 months of age (3, 5), neither the SKO nor *Fxr*^-/-^ mice exhibited liver tumors at 60 weeks old. This could be due to possible differences in gut microbiome in different animal facilities. Nonetheless, 100% of the DKO mice developed liver tumors by 60 weeks of age and RNA-seq demonstrated increased expression of tumorigenic myc-target and IL6_JAK_STAT3_signaling pathways in DKO mice compared to SKO mice. Consistent with increased cell proliferation, DKO mice had more liver Ki67+ cells and higher Cyclin D1 expression compared to SKO mice.

FXR null mice are widely used to study bile acid toxicity and cholestasis. As *Fxr*^-/-^ mice age to 24 weeks, hepatic XBP1 and other UPR pathways were activated due to presumed increased bile acid-induced ER stress. The inability to appropriately upregulate liver XBP1 resulted in increased liver injury, apoptosis, fibrosis and promoted spontaneous liver tumor development in *Fxr*^-/-^ mice. These data suggest that hepatic XBP1 expression is protective in *Fxr*^-/-^ mice, which may be due to a reduction of ER stress and/or crosstalk between liver XBP1 and FXR signaling. This may have important implications on the pathogenesis and treatment of cholestatic liver diseases.

## Acknowledgement

The authors wish to thank the Northwestern University Center of Genetic Medicine NUSeq core facility for performing RNA-seq and the Mouse Histology and Phenotyping Laboratory for performing the histologic staining.

