## Supplementary Figure Legends for "Hepatic Deletion of X-box Binding Protein 1 in Farnesoid X Receptor Null Mice Leads to Enhanced Liver Injury"

**Supplementary Figure S1: Hepatic expression of the IRE1α/XBP1 pathway proteins in WT and *Fxr*^-/-^ mice.** (A) Densitometry analysis of nuclear XBP1s, whole liver homogenate p-IRE1α and IRE1α protein expression normalized to loading control in 10-week-old (n=6) and 24-week-old (n=5) male WT and *Fxr*^-/-^ (KO) mice. *P<0.05, ***P<0.001. (B) Hepatic *Fxr* gene expression analysis by qPCR confirming the FXR phenotype of the *Fxr*^-/-^ mice. ****P<0.0001.

**Supplementary Figure S2: Hepatic expression of the PERK pathway proteins in WT and *Fxr*^-/-^ mice.** Densitometry analysis of the whole liver homogenate p-PERK, PERK, p-eIF2α, eIF2α, ATF4 and nuclear CHOP protein expression normalized to loading control in 10-week-old (n=6) and 24-week-old (n=5) male WT and *Fxr*^-/-^ (KO) mice. *P<0.05, **P<0.01, ***P<0.001, ****P<0.0001.

**Supplementary Figure S3: Confirmation of the FXR and XBP1 phenotypes of *Fxr*^-/-^*Xbp1*^fl/fl^** **and *Fxr*^-/-^*Xbp1*^LKO^ mice.** Hepatic gene expression of *Xbp1* and *Fxr* was measured by qPCR in male chow-fed 10-week-old *Fxr*^-/-^*Xbp1*^fl/fl^ (SKO, n=4) and *Fxr*^-/-^*Xbp1*^LKO^ mice (DKO, n=6). WT mice was used as a positive control for *Fxr* expression. ****P<0.0001.

**Supplementary Figure S4: Hepatic expression of the PERK pathway proteins in *Fxr*^-/-^*Xbp1*^fl/fl^** **and *Fxr*^-/-^*Xbp1*^LKO^ mice.** Liver protein was isolated from 10-week-old male *Fxr*^-/-^*Xbp1*^fl/fl^ (SKO, n=4) and *Fxr*^-/-^*Xbp1*^LKO^ mice (DKO, n=6). (A) Western blotting examining p-PERK, PERK, p-eIF2α, eIF2α, and ATF4 in whole liver. GAPDH was used as a loading control. (B) Densitometry analysis of the protein expression normalized to GAPDH. *P<0.05, **P<0.01.

**Supplementary Figure S5: Hepatic expression of Cyclin D1 in *Fxr*^-/-^*Xbp1*^fl/fl^**, ***Fxr*^-/-^*Xbp1*^LKO^, *Xbp1*^fl/fl^ and *Xbp1*^LKO^ mice.** Western blotting demonstrating hepatic Cyclin D1 protein expression in (A) 10-week-old male *Fxr*^-/-^*Xbp1*^fl/fl^ (SKO, n=4) and *Fxr*^-/-^*Xbp1*^LKO^ mice (DKO, n=6); (B) 24-week-old male SKO (n=4) and DKO (n=6) mice; (C) 40-week-old male SKO (n=5) and DKO (n=6) mice; and (D)10-week-old male *Xbp1*^fl/fl^ and *Xbp1*^LKO^ mice (n=4 per group). GAPDH was used as a loading control.
