## Supplementary figures and images for "Hepatic Deletion of X-box Binding Protein 1 in Farnesoid X Receptor Null Mice Leads to Enhanced Liver Injury"

### Supplementary Figure S1

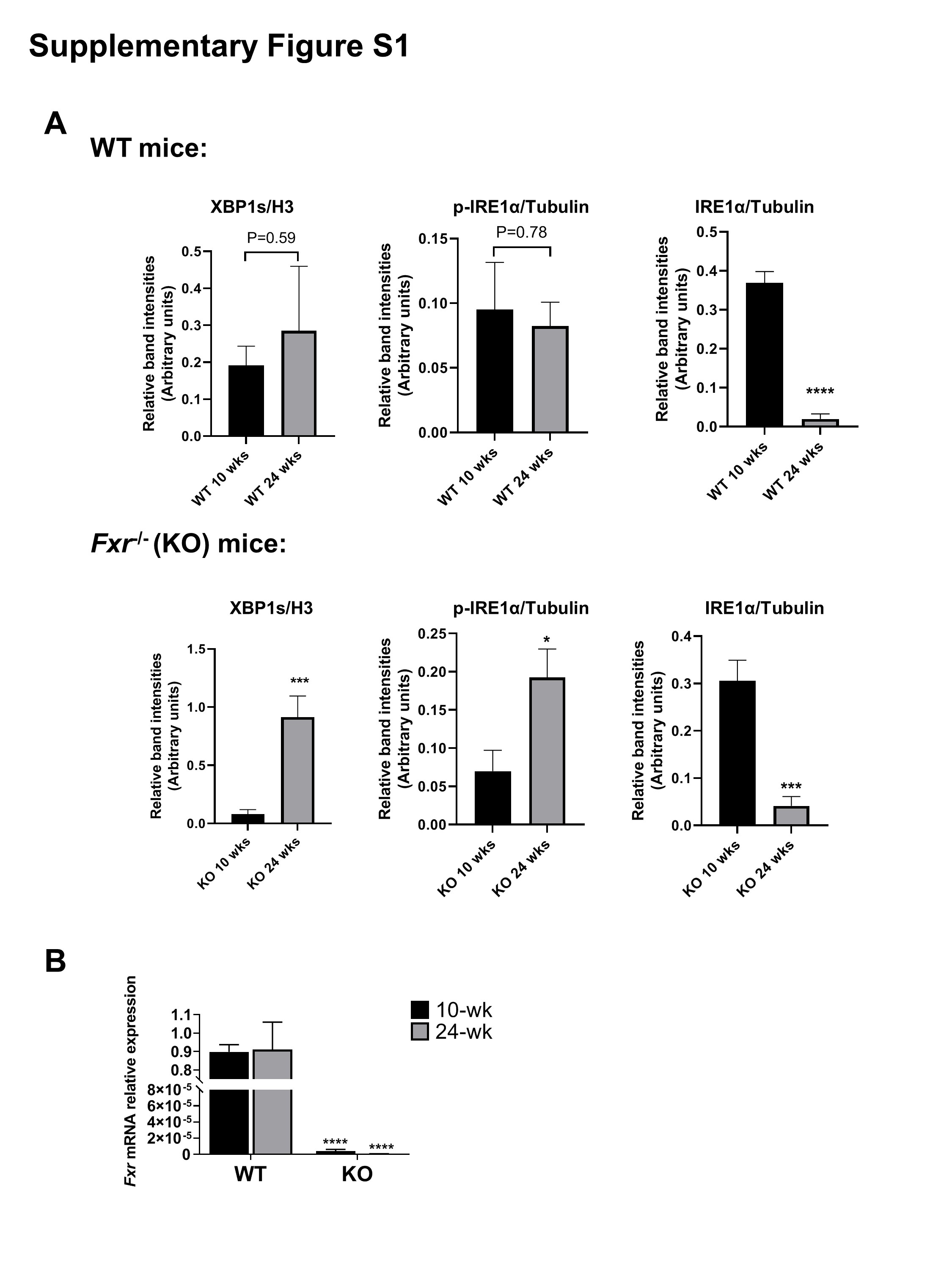

### Supplementary Figure S2

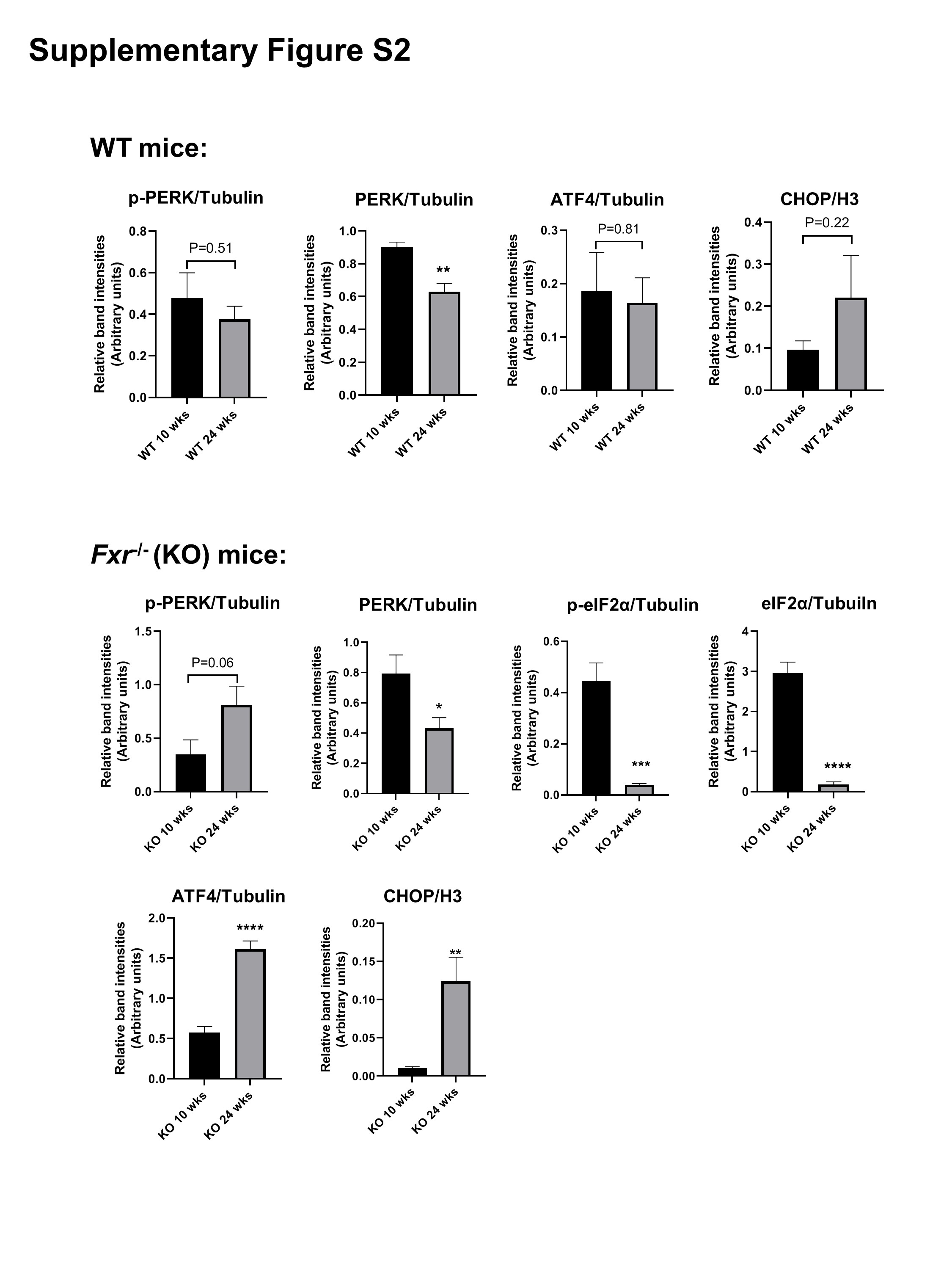

### Supplementary Figure S3

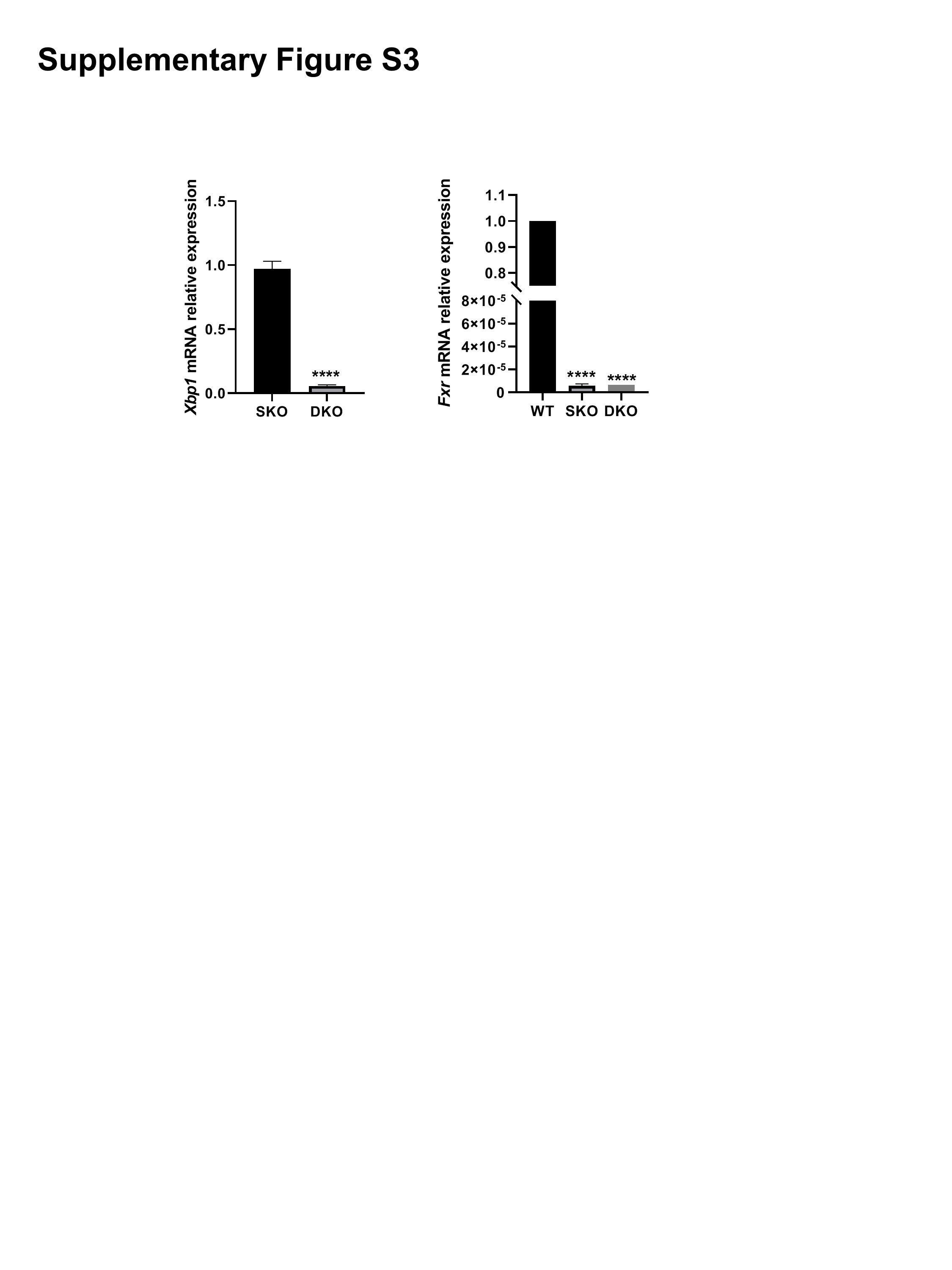

### Supplementary Figure S4

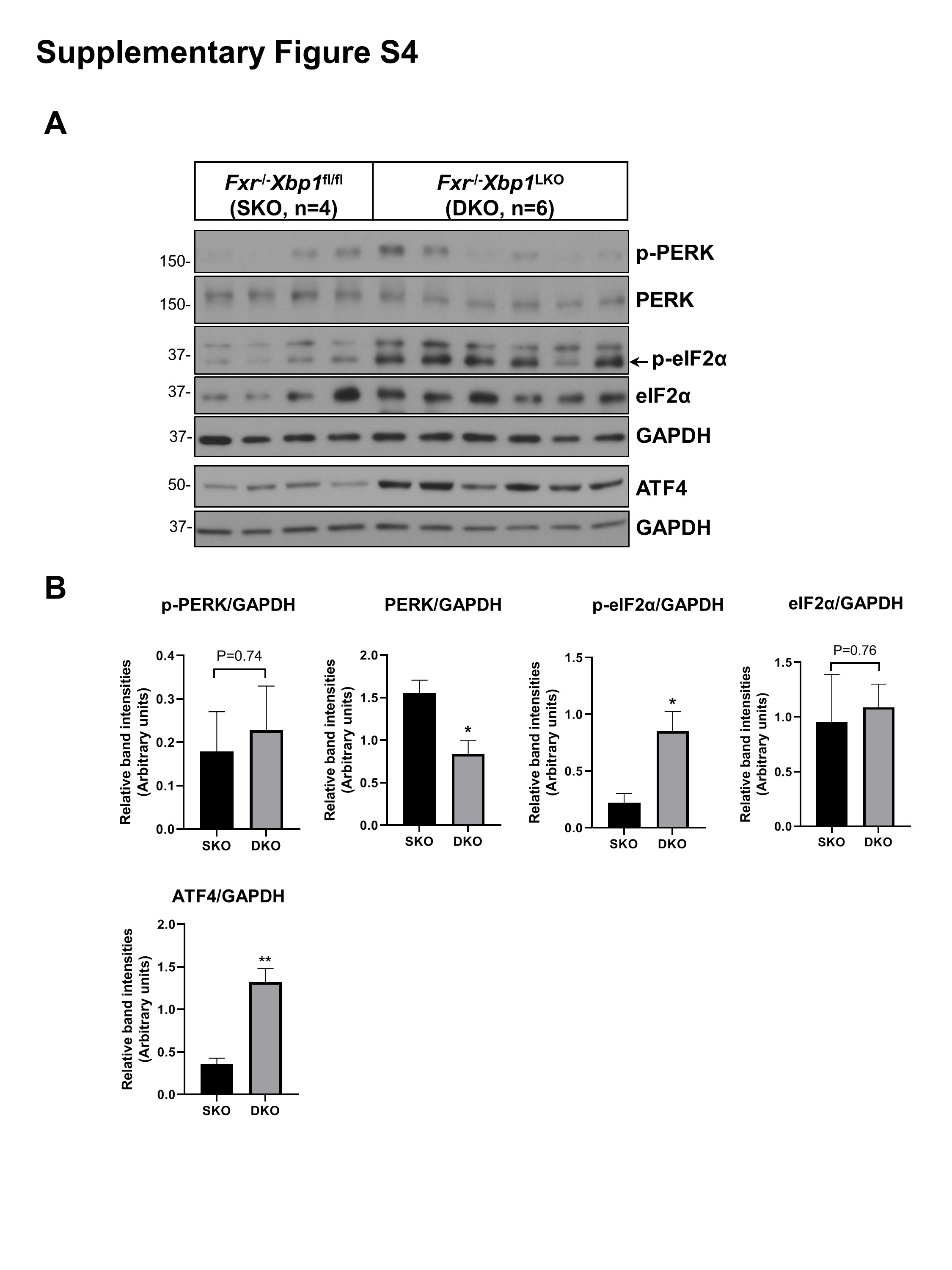

### Supplementary Figure S5

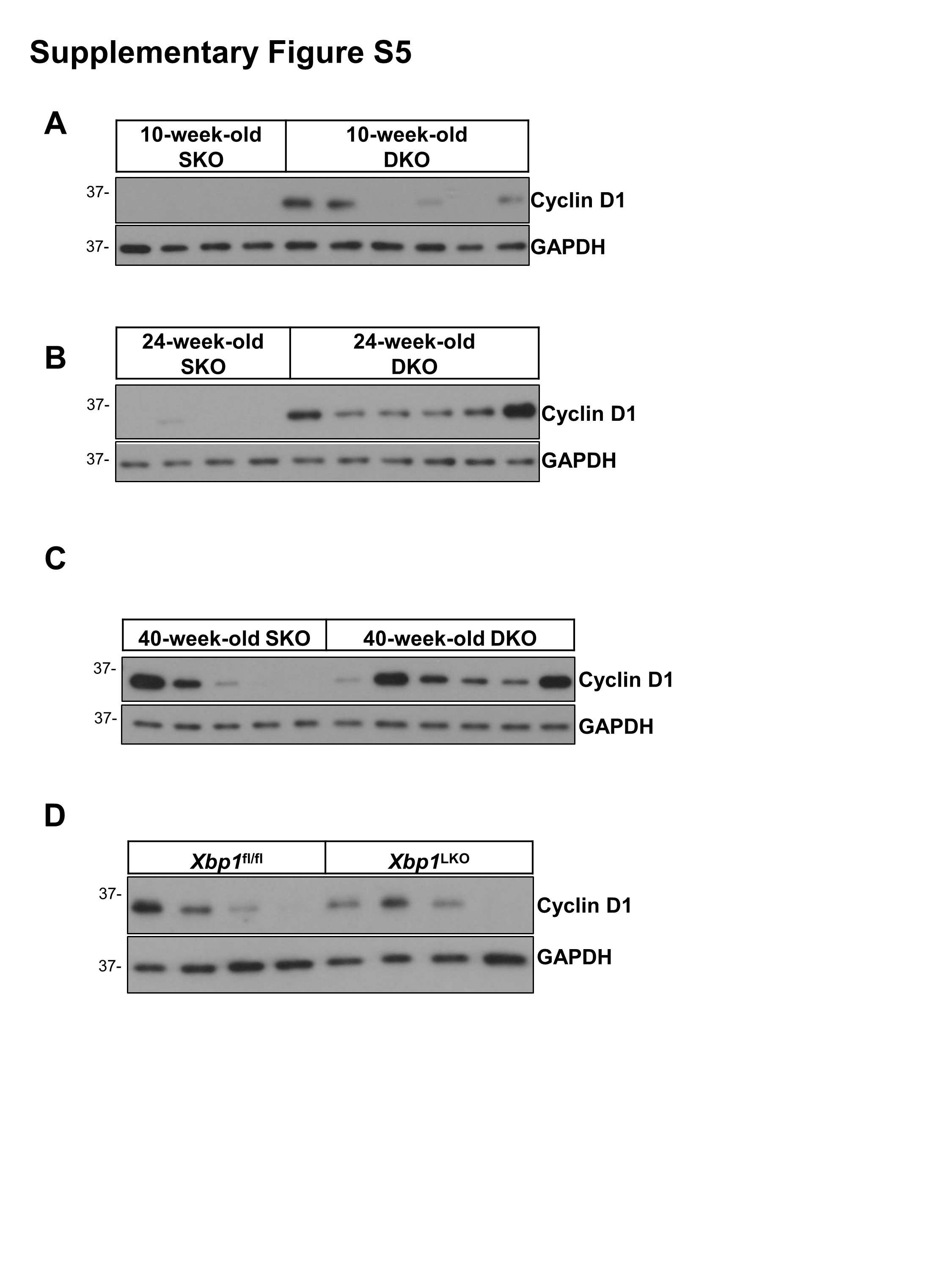
